# Phylogenomic analysis of the APETALA2 transcription factor subfamily across angiosperms reveals both deep conservation and lineage-specific patterns

**DOI:** 10.1101/859926

**Authors:** Merijn H.L. Kerstens, M. Eric Schranz, Klaas Bouwmeester

## Abstract

The APETALA2 (AP2) subfamily of transcription factors are key regulators of angiosperm root, shoot, flower, and embryo development. The broad diversity of anatomical and morphological structures is potentially associated with the genomic dynamics of the *AP2* subfamily. However, a comprehensive phylogenomic analysis of the *AP2* subfamily across angiosperms is lacking. We combined phylogenetic and synteny analysis of distinct *AP2* subclades in the completed genomes of 107 angiosperm species. We identified major changes in copy number variation and genomic context within subclades across lineages, and discuss how these changes may have contributed to the evolution of lineage-specific traits. Multiple *AP2* subclades show highly conserved patterns of copy number and synteny across angiosperms, while others are more dynamic and show distinct lineage-specific patterns. As examples of lineage-specific morphological divergence due to *AP2* subclade dynamics, we hypothesize that that loss of *PLETHORA1/2* in monocots correlates with the absence of taproots, whereas independent lineage-specific changes of *PLETHORA4*/*BABY BOOM* and *WRINKLED1* genes in Brassicaceae and monocots point towards regulatory divergence of embryogenesis between these lineages. Additionally, copy number expansion of *TOE1* and *TOE3/AP2* in asterids is implicated with differential regulation of flower development. Moreover, we show that the genomic context of *AP2*s is in general highly specialised per angiosperm lineage. Our study is the first to shed light on the evolutionary divergence of the *AP2* subfamily subclades across major angiosperm lineages and emphasises the need for lineage-specific characterisation of developmental networks to further understand trait variability.

**SIGNIFICANCE STATEMENT:** APETALA2 transcription factors are crucial regulators of embryogenesis and post-embryonic development in plants. Characterising the genomic dynamics of *APETALA2* genes across 107 angiosperms provided evolutionary insights into gene-family and morphological divergence across major angiosperm lineages.

## INTRODUCTION

Angiosperms display an extraordinary diversity in anatomy and morphology across a large number of phylogenetic lineages, each with their own characteristic traits (Stevens, 2001; Leebens-Mack *et al*., 2019). The most distinctive lineages are the monocots and the eudicots, which can be characterized by having either one or two cotyledons, respectively. Many crop and model species are distributed across the phylogenetic tree, such as in the rosids (i.e. soybean, strawberry, cassava, cucumber, *Arabidopsis*, cotton and orange), the asterids (tomato, coffee, carrot and lettuce) and in the monocots (wheat, corn, pineapple, banana, oil palm). Phenotypic diversity in angiosperm lineages can be largely explained by patterns of gene diversification. A major driver of genetic diversification in plants is gene duplication, which promotes functional redundancy leading to the development of novel traits (Van de Peer *et al.*, 2009; Guo, 2013; Panchy *et al.*, 2016). Various processes are thought to facilitate gene duplication, of which polyploidisation events are regarded as having the most profound impact (Panchy *et al.*, 2016). Polyploidisation is often followed by the process of diploidisation, which reshuffles genome structure by chromosomal rearrangements and massive gene loss with many duplicated genes returning to a single-copy state (Dodsworth *et al.*, 2016). On a more local scale, numerous genes undergo a process of constant gene birth-death by tandem duplication, transposition and retroduplication (Panchy *et al.*, 2016). Both global and local duplication mechanisms provide the raw material needed for gene diversification, thereby affecting angiosperm trait evolution.

Retention of genomic position and gene order over the course of evolution, called synteny, is thought to be indicative of conservation of gene function (Dewey, 2011; Lv *et al.*, 2011). Hence, studying the evolutionary history of gene synteny in related clades can reveal key events in the acquisition of novel traits caused by gene duplications and rearrangement of genomic context (Tang *et al.*, 2008; Dewey, 2011; Jiao and Paterson, 2014). Recently, we developed a pipeline that exploits network clustering to study syntenic relationships between genes, overcoming challenges imposed by pairwise interspecies comparisons (Zhao and Schranz, 2017). With this approach, new evolutionary trends could be inferred for MADS-box and *LEA* gene families across 51 plant species (Zhao *et al.*, 2017; Artur *et al.*, 2019). More recently, this approach was used to examine overall syntenic properties and genomic differences between angiosperms and mammals (Zhao and Schranz, 2019). These studies demonstrate the potential of network clustering to simultaneously study synteny of diverse gene families in multiple species.

The *APETALA2/ETHYLENE-RESPONSIVE ELEMENT BINDING PROTEIN* (*AP2/EREBP*) superfamily is one of the most prominent transcription factor families regulating plant development and stress responses (Riechmann and Meyerowitz, 1998). It can be divided into two main subfamilies: the *AP2* subfamily, whose members contain two AP2 domains, and the *EREBP* subfamily containing a single AP2 domain (Riechmann and Meyerowitz, 1998). The *AP2* subfamily can be further divided in the *euANT, basalANT* and *euAP2* clades, of which members have been demonstrated in *Arabidopsis thaliana* to play key roles in diverse developmental processes (Kim *et al.*, 2006). The *euANT* gene *AINTEGUMENTA* (*ANT*) was shown to regulate floral organ identity and shoot meristem maintenance (Elliott *et al.*, 1996; Mudunkothge and Krizek, 2012). The six *PLETHORA* genes (*PLT1, PLT2, PLT3/AIL6, PLT4/BABY BOOM, PLT5/AIL5, PLT7/AIL7*) are redundant regulators of multiple key processes, including root and shoot development, phyllotaxis/rhizotaxis and embryogenesis (Aida *et al.*, 2004; Galinha *et al.*, 2007; Smith and Long, 2010; Prasad *et al.*, 2011; Hofhuis *et al*. 2013, Horstman *et al.*, 2014). The *basalANT* genes *WRI1, WRI3* and *WRI4* (*WRINKLED1*/*3*/*4*) govern fatty acid metabolism in the embryo and flower (Cernac and Benning, 2004; To *et al.*, 2012). Within the *euAP2* clade, *AP2* controls flower development and floral organ identity (Jofuku *et al.*, 1994; Okamuro *et al.*, 1997), while *TOE1* and *TOE3* (*TARGET OF EAT1/3*) regulate flowering (Zhu and Helliwell, 2011; Zhang *et al.*, 2015).

Having pivotal regulatory roles in multiple developmental processes, genome dynamics within the *AP2* subfamily can have profound effects on the morphological characteristics of angiosperms. Thus far, the *AP2* subfamily has only been studied in the context of a subset of subclades, a small number of species, or at low resolution in combination with the *EREBP* subfamily (Rashid *et al.*, 2012; Song *et al.*, 2013; Lakhwani *et al.*, 2016; Zumajo-Cardona and Pabón-Mora, 2016; Najafi *et al.*, 2018; Li *et al.*, 2018; Leebens-Mack *et al*., 2019; Wang *et al.*, 2019). To elucidate the evolution of the *AP2* subfamily in greater detail, it is important to assess all subclades of this subfamily in an angiosperm-wide manner.

Here, we determined the dynamics in copy number variation and syntenic conservation of *AP2* subfamily genes across all major angiosperm lineages. By combining phylogenetics and synteny network clustering, we identified distinct genomic patterns in the *AP2* subfamily, uncovering evolutionary forces that drive morphological and developmental diversity in angiosperms.

## RESULTS AND DISCUSSION

### Synteny conservation in *AP2* subclades of angiosperms

Phylogenomic analysis of the *AP2* subfamily was conducted by combining conventional phylogenetic and syntelog clustering methods (Fig. 1B, Zhao and Schranz, 2019). The annotated proteins of 107 angiosperm species, including two basal eudicots, were mined for protein sequences containing two repeated AP2 domains using HMMER (Fig. 1B). The resulting hits were termed ‘HMMERlogs’, homologous proteins that share identical domain compositions (Fig. 1A). In total, we identified 2171 AP2 HMMERlogs across 107 angiosperm proteomes (Table S1). Trimmed and aligned HMMERlog sequences were used for phylogenetic gene-tree analysis. In parallel, we performed synteny network analyses to identify *AP2* genes localised in regions of similar genomic context, termed ‘syntelogs’ (Fig. 1A). Out of the 2171 AP2 HMMERlogs, 1570 are connected by synteny to at least one other HMMERlog (Table S1).

**Figure 1.**
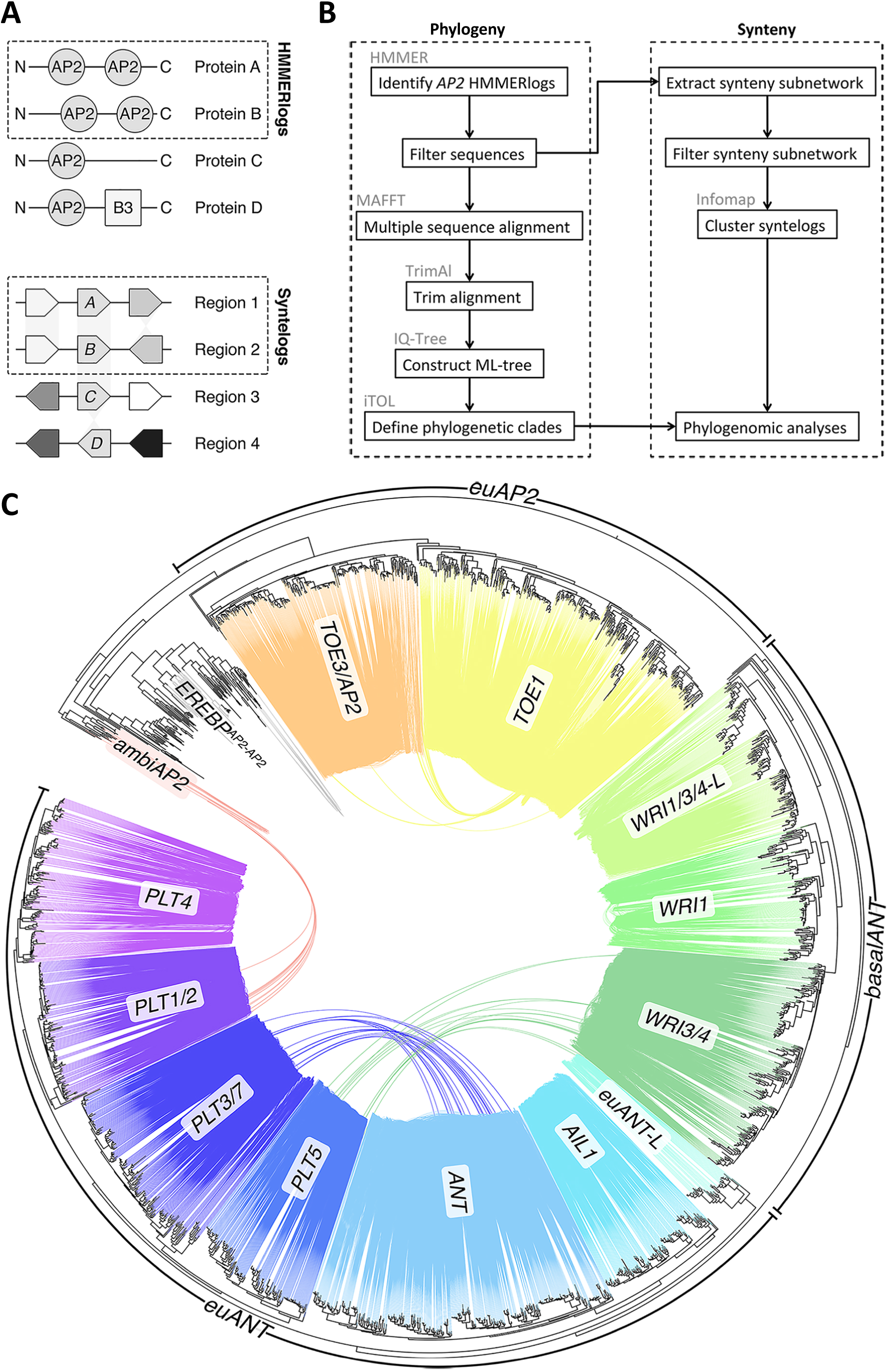
Phylogenetic and syntenic relationships of the *AP2* subfamily in angiosperms. (**A**) Schematic drawing illustrating the terms ‘HMMERlog’ (upper panel) and ‘syntelog’ (lower panel). HMMERlogs are defined as homologs with an identical domain composition (proteins A and B, both having two AP2 domains). Syntelogs are genes sharing conserved syntenic regions (e.g. genes *A* and *B* in Regions 1 and 2) (**B**) Bioinformatics workflow for HMMERlog identification and phylogenomic analyses. Main analysis steps and used software tools are indicated. (**C**) Phylogenetic and synteny network analysis of AP2 HMMERlogs in 107 angiosperm genomes. Coloured lines indicate strong conservation of synteny between gene pairs within and between AP2 subclades. Weaker syntenic connections between subclades are shown in grey in Fig. S1. Subclades are named following the *Arabidopsis* gene nomenclature. Subclades lacking *Arabidopsis* orthologs are annotated with the suffix -L (-like). The phylogenetic tree was inferred by ML analysis using 1000 bootstraps.

To gain insight into conservation and divergence of genomic context across the *AP2* subfamily of angiosperms, syntenic connections were plotted on the phylogenetic tree (Fig. 1; S1). Within the phylogenetic tree, three major clades and 14 distinct *AP2* subclades can be distinguished (Fig. 1C). Each *AP2* subclade consists of highly interconnected HMMERlogs. This syntenic signal is in strong congruence with the observed phylogeny (Fig. 1C), and supports the described gene phylogeny of the *AP2* subfamily in angiosperms. An exception is the *EREBP*^*AP2-AP2*^ subclade consisting of HMMERlogs with two AP2 domains, but having overall higher homology with EREBP proteins. HMMERlogs of this subclade were not included in further analyses. One group of 17 AP2 HMMERlogs belongs to neither the *euAP2* nor *basalANT* clade, and was termed *‘ambiAP2’* for its ambiguity. Strong syntenic connections are found between HMMERlogs of four subclade pairs: subclades *ANT* and *PLT3/7, PLT1/2* and *ambiAP2, PLT5* and *WRI3/4*, and *TOE3/AP2* and *TOE1* (Fig. 1C), respectively. Two of these subclade pairs contribute to the same plant developmental processes in *Arabidopsis. ANT, PLT3* and *PLT7* play key roles in shoot meristem maintenance (Mudunkothge and Krizek, 2012), whereas *AP2, TOE1* and *TOE3* regulate flowering (Jofuku *et al.*, 1994; Okamuro *et al.*, 1997; Zhu and Helliwell, 2011; Zhang *et al.*, 2015). Similar patterns of shared synteny were detected for the Type II MADS-box subclade pairs *AGL6*-*TM3, SEP1*-*SQUA* and *SEP3-FLC* (Ruelens *et al.* 2013; Zhao *et al*. 2017). This suggests that the functions of these AP2 HMMERlogs are facilitated by a shared genomic context. As of yet, a potential shared function between the other two subclade pairs remains unclear.

### *AP2* copy number variation as potential driver of morphological diversity in angiosperms

We explored copy number variation in *AP2* subclades to further understand morphological divergence in angiosperms (Fig. 2). Across most angiosperms, the *PLT1/2* subclade displayed relatively little variation in copy number. However, all monocots lack HMMERlogs of this subclade, suggesting gene loss during early monocot evolution (Fig. 2). Since *Arabidopsis PLT1* and *PLT2* are crucial regulators of root development (Aida *et al.*, 2004; Galinha *et al.*, 2007), this process is potentially regulated in a different fashion in monocots. In support of this, the basal eudicot *Amborella trichopoda* contains a single PLT1/2 HMMERlog and has a taproot system (Trueba *et al.*, 2016). Most eudicots have a taproot system; a primary root that develops directly from an embryonic root, out of which secondary roots emerge. In monocots, embryonic roots are aborted, inducing adventitious root formation that leads to the formation of a fibrous root system. While monocots lack PLT1/2 HMMERlogs, they do contain monocot-specific *euANT-L* HMMERlogs that potentially assume the role of *PLT1/*2 in root development (Fig. 2). Likewise, since *Arabidopsis PLT4* also redundantly orchestrates root development (Galinha *et al.*, 2007), the increase in number of PLT4 HMMERlogs in monocots compared to eudicots could compensate for the loss of PLT1/2 genes. Regardless of their main root system, both eudicots and monocots form secondary roots. In *Arabidopsis, PLT3, PLT5* and *PLT7* govern rhizotaxis (Hofhuis *et al.*, 2013). For the *PLT5* subclade, the number of copies is relatively constant across angiosperms, also between eudicots and monocots, although the latter lineage most often has two copies instead of one. Copy number variation of the *PLT3/7* subclade is also low, whilst the increased copy number in Brassicaceae is potentially caused by recent whole-genome duplications. The relative conservation of these two subclades might indicate that the secondary root developmental programme is conserved between eudicots and monocots, despite distinct differences in their main root system. Moreover, since *PLT3, PLT5* and *PLT7* also play key roles in *Arabidopsis* phyllotaxis (Prasad *et al.*, 2011), our findings suggest that lateral organ positioning in general is conserved between these lineages.

**Figure 2.**
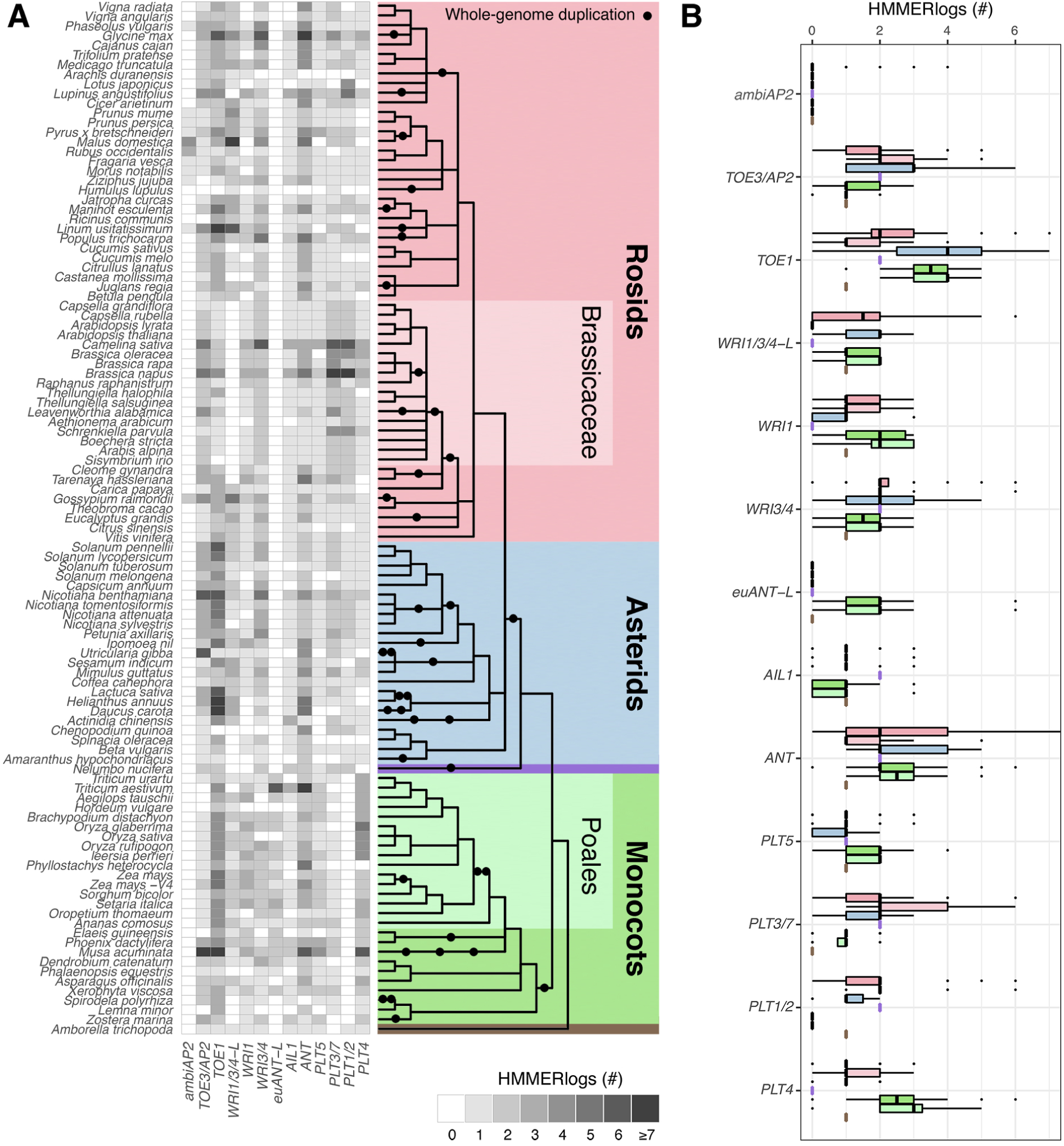
Dynamic copy number variation within the angiosperm *AP2* subfamily (**A**) Copy number variation of AP2 HMMERlogs in angiosperms. Cell shading indicates the number of HMMERlogs across species. Colours represent angiosperm taxa; i.e. rosids (pink), asterids (blue), monocots (green), Brassicaceae (light pink), Poales (light green), and the basal eudicots *Nelumbo nucifera* (purple) and *Amborella trichopoda* (brown), respectively. Whole-genome duplication events are indicated by black dots based on recent studies (Barker *et al.*, 2016; Zhao *et al*., 2017, Clark and Donoghue, 2018; Leebens-Mack *et al*., 2019; Qiao *et al.*, 2019). (**B**) Box plot showing the distribution of AP2 HMMERlogs per subclade across angiosperm lineages. Colours are defined as in **A**.

Monocot and eudicot embryogenesis differ in the number of developed cotyledons, indicating that their embryo developmental programmes are likely regulated in a very different fashion. The question is thus raised, how such differences are reflected by copy number variation in the *AP2* subfamily. The absence of both *PLT2* and *PLT4* was shown to be embryo lethal in *Arabidopsis* (Galinha *et al.*, 2007). As already pointed out, monocots lack PLT1/2 HMMERlogs, but have more *PLT4* copies (Fig. 2). This potentially buffers the loss of PLT1/2 function. Alternatively, monocot-specific euANT-L HMMERlogs may have defining roles during embryogenesis.

The *WRI1, WRI3/4* and *WRI1/3/4-L* subclades, consisting of transcriptional regulators of fatty acid synthesis (Cernac and Benning, 2004; To *et al.*, 2012), are largely conserved in copy number across angiosperms. Brassicaceae, however, lack *WRI1/3/4-L* orthologs, suggesting an evolutionary loss of a distinct subclade of fatty acids regulators.

Floral morphology in angiosperms is extremely diverse, as specified through the modular ‘ABC’ model. In comparison to other angiosperm lineages, asterids exhibit duplicate retention of a large number of TOE1 HMMERlogs, and TOE3/AP2 HMMERlogs to a lesser extent. Potentially, this is linked to a divergent form of modular flower development in asterids. In support of this, major differences in regulation of floral organ patterning were found between *Arabidopsis* and *Petunia hybrida*, belonging to the rosid and asterid lineages, respectively. In *Arabidopsis*, the transcription factor AP2 has an A-class function and antagonizes B- and C-class genes (Krogan *et al.*, 2012). However, the closest homologs of *AP2* in *Petunia* – i.e. *ROB1, ROB2* and *ROB3* (*REPRESSOR OF B-FUNCTION1/2/3*), found in the *TOE3/AP2* subclade - only antagonize B-class genes (Morel *et al.*, 2017). The *Petunia BEN* (*BLIND ENHANCER*) gene has an identical role to *Arabidopsis AP2* by repressing both B- and C-class genes, but instead belongs to the *TOE1* subclade. Thus, whereas *Arabidopsis TOE1* is a repressor of flowering time but does not regulate flower patterning, *BEN* acts in flower patterning but does not affect flowering time (Morel *et al.*, 2017). In addition, *BEN* and *ROB* genes also regulate nectary size (Morel *et al.*, 2018). This points out that expansion of the *euAP2* clade in asterids has led to more complex patterning during flower development.

The number of ANT HMMERlogs is particularly variable across rosids. For example, *ANT* copy number is high in Fabaceae and Cucurbitaceae (3-4) and low in Brassicaceae (∼1). Although we are unable to pinpoint a specific trait that correlates with this, the defining role of *ANT* in cell proliferation in seeds, leaves and flowers (Krizek, 1999; Mizukami and Fischer, 2000; Confalonieri *et al.*, 2014) suggests that *ANT* copy number might very well be involved in shaping the distinctive morphology of these structures in different angiosperm lineages. Likewise, ambiAP2 HMMERlogs are present almost exclusively in the Rosaceae, but without knowledge of their function, we could not identify a clear link to developmental traits.

### The angiosperm *AP2* subfamily is impacted by ancestral and lineage-specific synteny

Extensive synteny conservation is often linked with preservation of gene function (Dewey, 2011; Lv *et al.*, 2011). As such, we investigated synteny variation within *AP2* subclades across angiosperms. For each angiosperm species, we determined copy number variation within each syntelog cluster and performed hierarchical clustering (Fig. 3A). In this way, 27 syntelog clusters were identified, including all subclades except *ambiAP2*, each representing a distinct genomic context (Fig. 3B). Our data suggest that the genomic context of many *AP2* subclades, despite their conserved roles in crucial developmental processes, varies across angiosperms. This demonstrates that synteny in the *AP2* subfamily is largely lineage-specific, with only few subclades being deeply conserved across all angiosperms (Fig. 3). The level of synteny conservation does not correlate with the number of HMMERlogs in a subclade. For example, the *ANT* subclade is the second largest in number of HMMERlogs, but only represented in a single syntelog cluster. This reassures that variation in syntelog cluster size is not merely a product of the number of HMMERlog within a subclade.

**Figure 3.**
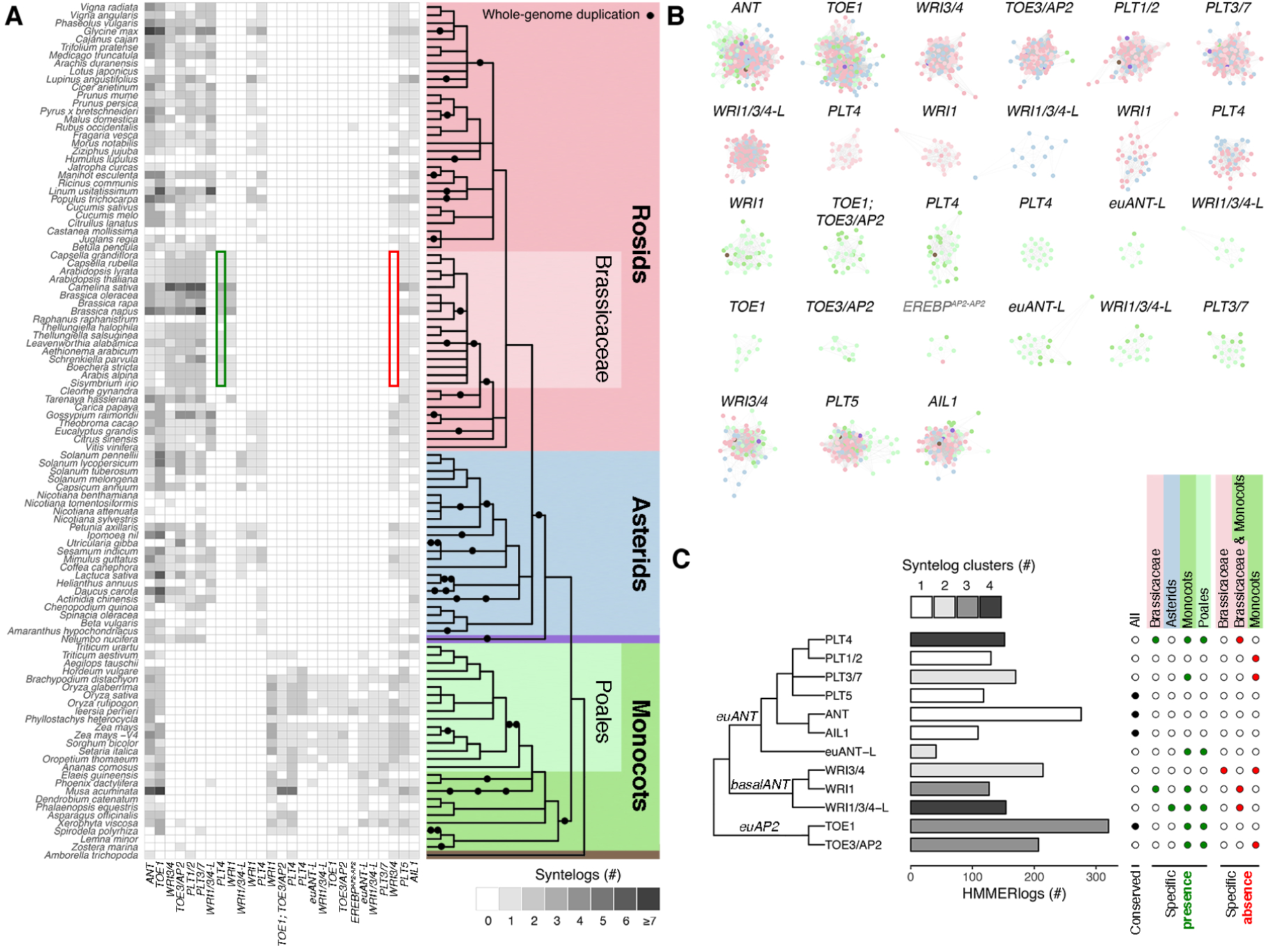
Ancestral and lineage-specific synteny in the angiosperm *AP2* family. (**A**) Phylogenomic profile of *AP2* syntelogs across angiosperm genomes. Columns display syntelog clusters. Cell shading indicates the number of syntelogs per species. Examples of lineage-specific synteny are indicated in green (presence) and red (absence), respectively. (**B**) Visual representation of syntelog clusters following the order of columns in **A** (left to right). Nodes indicate syntelogs, coloured according to the angiosperm phylogeny. Edges display syntenic connections between syntelogs. Distances between nodes scales with the number of connections between nodes. (**C**) Bar plot displaying the number of syntelog clusters and total number of HMMERlogs per *AP2* subclade (left panel). The right panel displays conserved synteny (black dots) and presence/absence of lineage-specific synteny (green/red dots, respectively) across angiosperm lineages.

As a next step, we quantified the level of synteny conservation across angiosperms (Fig. 3C). Four *AP2* subclades, each consisting of a single syntelog cluster, display extremely strong conservation of synteny across all angiosperm species. This includes the *ANT, AIL1, PLT1/2* (although only present in eudicots) and *PLT5* subclades, and proposes the existence of strict positional constraints to conserve gene function. As such, genes belonging to these subclades are likely to share the same core functions across all angiosperms. More dynamic patterns of synteny can be observed in the other *AP2* subclades, each displaying specific presence or absence of synteny in different angiosperm lineages. For instance, within the *PLT3/7* subclade two separate genomic contexts can be observed, i.e. specific presence of synteny in monocots, and specific absence of synteny in eudicots. Since *PLT3* and *PLT7* play key roles in phyllotaxis and rhizotaxis in *Arabidopsis* (Prasad *et al*., 2011; Hofhuis *et al.*, 2013), our data suggest that these processes are differently regulated in monocots. The *PLT4* subclade exhibits markedly more lineage-specific synteny than its related subclades, including lineage-specificity in Brassicaceae, monocots and Poales. This potentially influences regulation of embryogenesis and root development in different angiosperm lineages. The *WRI* subclades are generally more dynamic with respect to their genomic position in Brassicaceae, asterids and monocots, suggesting differential regulation of fatty acid synthesis in these lineages.

*WRI1* and *PLT4* are the only subclades that exhibit Brassicaceae-specific synteny. *Arabidopsis* WRI1 and PLT4 are direct downstream targets of the LAFL (LEC1-ABI3-FUS3-LEC2) transcription factor network that regulates embryogenesis (Jia *et al.*, 2014; Horstman *et al.*, 2017). This finding highlights that Brassicaceae evolved unique mechanisms to regulate embryogenesis. This is reinforced by the fact that *Arabidopsis LEC1*, one of the four core *LAFL* genes, also belongs to a Brassicaceae-specific syntelog cluster (Fig. S2). The other three *LAFL* genes, however, do not exhibit Brassicaceae-specificity (Fig. S2). Future research will be needed to test the hypothesis that the unique syntenic positions of *WRI1, PLT4* and *LEC1* affect the promoter and/or chromatin dynamics of these genes and thus create a Brassicaceae-specific embryo development regulatory network.

Within the two *TOE* subclades, monocots and eudicots are in separate synteny clusters, implying regulation of distinct flower development in these major angiosperm lineages. These analyses are the first to define lineage-specific synteny relationships within the *AP2* subfamily, and form a framework to unravel differential transcriptional regulation of various developmental programmes in angiosperms.

### Synteny of the *AP2* subfamily is specialised per angiosperm lineage

The *AP2* subfamily displays a remarkably high level of lineage-specific synteny, especially since this gene family orchestrates multiple developmental programs in angiosperms. Therefore, we determined how this degree of lineage-specificity relates to the presumably more dynamic sister subfamily of EREBP transcription factors, which play essential roles in regulating responses to biotic and abiotic stress (Licausi *et al.*, 2013). In comparison to the *AP2* subfamily, *EREBP* genes are more strongly interconnected by synteny, even though the ratio of syntelogs to HMMERlogs is identical (Fig. 4A, left and middle panel). This is likely due to larger number of *EREBP* syntelogs per cluster (Fig. 4A, right panel). This implies that the *EREBP* subfamily has had more retention of duplicated genes or experienced duplication more often, but it does not reveal any information on the degree of lineage-specific synteny. Hence, we performed phylogenomic analyses of the *EREBP* subfamily and classified subclades as described by Nakano *et al.*, 2006 (Fig. S3-S6). Similar to the *AP2* subfamily, *EREBP* subclades demonstrated copy number variation and lineage-specific synteny (Fig. S4-S6). To quantify this, we calculated a cluster overlap index of all pairwise interspecies comparisons. These cluster overlap indices can be used to compare relative conservation of synteny between the *AP2* and *EREBP* subfamilies (Fig. 4B). Of both related families, synteny is less conserved across all angiosperms in the *AP2* family. However, when the overall cluster overlap index is split into the five defined angiosperm lineages, conservation of *AP2* subfamily synteny is stronger than conservation of *EREBP* subfamily synteny in Brassicaceae, monocots and Poales. This suggests that the *AP2* genomic context is more specialized per lineage than that of the *EREBPs*.

**Figure 4.**
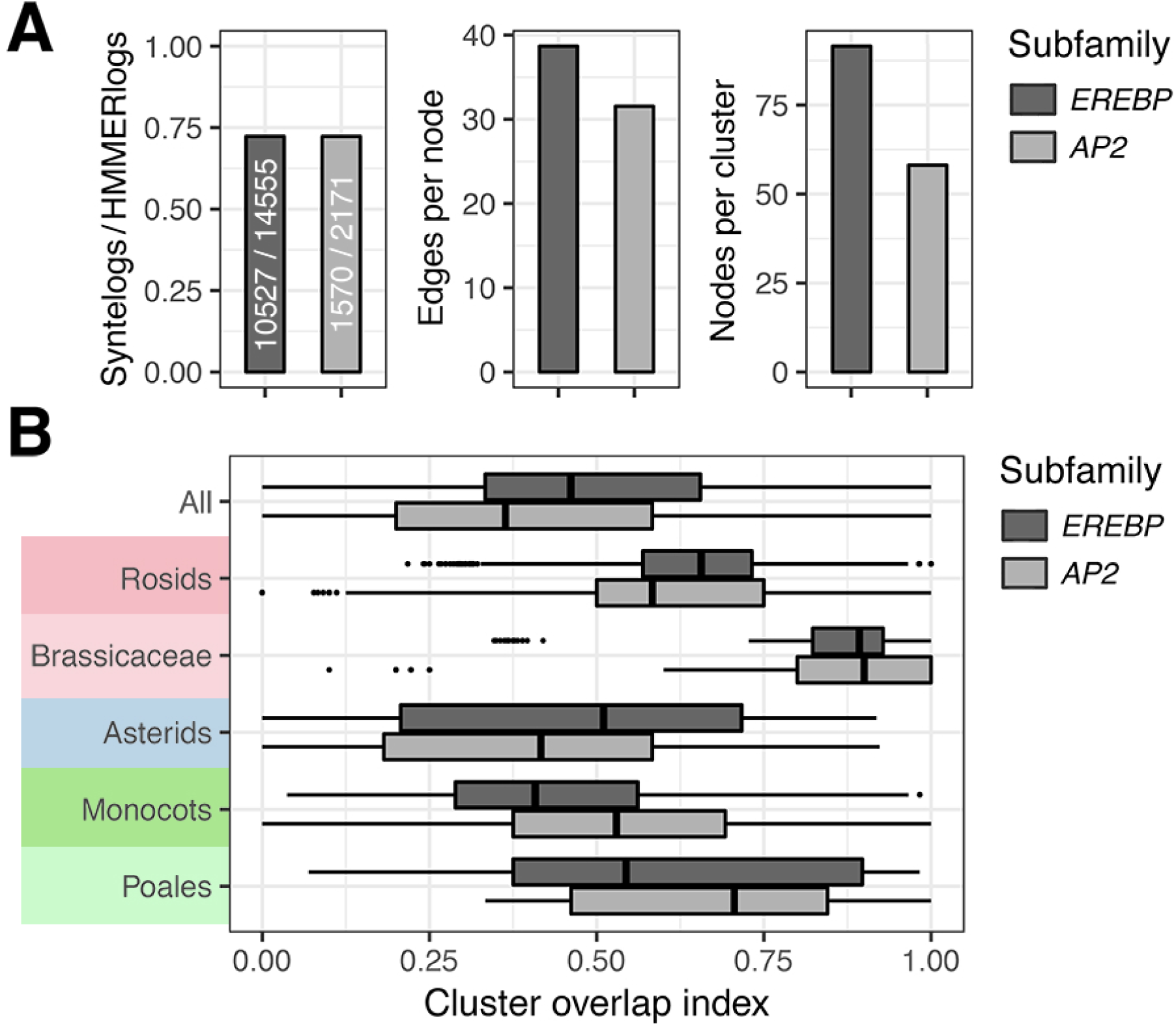
Synteny conservation in *AP2/EREBP* subfamilies varies per angiosperm lineage (**A**) Synteny network characteristics of the *AP2* and *EREBP* subfamilies. The ratio indicates the number of HMMERlogs with at least one syntenic connection. Nodes per cluster reflect syntelog cluster size. Edges per node reflect overall genomic connectedness. (**B**) Cluster overlap index of the *AP2* and *EREBP* syntelog networks per angiosperm lineage. Scale values range from 0 to 1, and display the proportion of HMMERlogs belonging to identical syntelog clusters.

## OUTLOOK

Our study exemplifies that combining phylogenetics and synteny networking (Fig. 1) is a powerful tool to investigate gene families, and the individual subclades therein, in large sets of species. Here, we provide two lines of potential evidence that associate variations in the *AP2* subfamily with angiosperm evolution. First, we reveal that angiosperm lineages differ in regard to gene copy number of phylogenetically distinct *AP2* subclades (Fig. 2). Secondly, we demonstrate that these subclades exhibit both deeply conserved and lineage-specific synteny (Fig. 3). Additionally, we show that the genomic contexts of *AP2* genes are specialized per angiosperm lineage (Fig. 4).

Through both a copy number and synteny perspective, we put forth a set of hypotheses that could be interesting to pursue in further research on the *AP2* subfamily. In the *PLT1/2* subclade, eudicots possess in general only one or two copies (Fig. 2B), all localised in a single genomic context (Fig. 3C). A similar observation is made for the *PLT5* and *AIL1* subclades across all angiosperms. Therefore, it seems that genes in these subclades experience a tight regulation of gene dosage and transcription. Overexpression of *Arabidopsis PLT5* was shown to induce formation of somatic embryos and enlargement of floral organs (Nole-Wilson *et al.*, 2005; Tsuwamoto *et al.*, 2010). Similarly, *Arabidopsis plt1,plt2* double mutants are disrupted in root meristem identity and develop short roots, whereas *PLT2* overexpression induces ectopic formation of root meristems (Aida *et al.*, 2004; Galinha *et al.*, 2007). No function has yet been attributed to *AIL1*. The extreme conservation of the *AIL1* subclade creates an extra incentive to functionally characterise its presumptive key role in plant development.

For other subclades, variation in copy number does not reflect variation in synteny. *ANT* genes vary largely in copy number, but only belong to a single syntelog cluster (Figs. 2B, 3C). This can be explained in two ways: i) flexibility in *ANT* gene dosage, but restriction in regulation, or ii) sub-functionalisation of *ANT* genes restricting variation in expression. The other way around, WRI1 HMMERlog copy number is more or less constant across different angiosperm lineages (Fig. 2B), but *WRI1* synteny is variable (Fig. 3C). Copy number variation in the *PLT3/7* subclade does not differ substantially between monocots and eudicots (Fig. 2B). The genomic context of *PLT3/7* subclade is however unique between both lineages (Fig. 3C), suggesting an evolutionary divergence in regulatory networks of phyllotaxis and rhizotaxis. In contrast, the *PLT5* subclade that also contributes to phyllotaxis and rhizotaxis (Prasad *et al*., 2011; Hofhuis *et al*., 2013) is strongly conserved (Fig. 3C). This suggests that lateral organ positioning has an ancestral component through *PLT5*, and a eudicot/monocot-specific regulatory component through *PLT3/7*. In conclusion, we show that combining phylogenetic analysis and syntelog clustering is a powerful tool to assess conservation and lineage-specificity of individual subclades within angiosperm gene families. This serves as a valuable resource for linking trait evolution to specific genomic events.

## EXPERIMENTAL PROCEDURES

### Identification of HMMERlogs in 107 angiosperm species

Proteomes of 107 angiosperm species were retrieved from public repositories as described by Zhao and Schranz (2019). Sequence similarity searches were performed using the AP2 alignment in Stockholm format (AP2, 2x PF00847; EREBP, 1x PF00847) using HMMER v.3.2.1. (El-Gebali *et al.*, 2019). Hits below the default inclusion threshold (E-value <0.01) were extracted. Protein sequences containing two AP2 domains were classified as AP2 subfamily members, and those with a single AP2 domain as EREBP subfamily members. Note that proteins with more than two AP2 domains were also assigned as AP2 members to account for erroneously recognised repeated domains. Protein sequences lacking a start codon were filtered out (Table S1). HMMERlogs containing NFYB domains (PF00808) or single B3 domains (PF02362) were identified in a similar way as described above.

### Multiple sequence alignment and phylogenetic analysis

Full-length protein sequences were aligned with MAFFT v7 using the FFT-NS-2 progressive algorithm with a gap penalty of 1.0 (Katoh *et al.*, 2017). Spuriously-sized sequences were filtered by length range (AP2, 200-800 amino acids; EREBP, 100-525; NFYB 100-350; and B3, 100-1250, respectively). Gapped positions in filtered multiple sequence alignments were removed by trimAl (Capella-Gutiérrez *et al.*, 2009). For AP2 and EREBP sequences, the automatic ‘-gappyout’ mode was used, which retained several hundred positions in the multiple sequence alignments including the AP2 domain(s). This setting proved to be unsuitable for trimming the more variable NFYB and B3 proteins. Instead, approximately 300 positions with the least gaps were kept by altering the ‘-gt’ and ‘-cons’ flags. Maximum-likelihood trees were constructed from trimmed alignments with IQ-TREE 1.6.10, using the LG substitution matrix and 1000 ultrafast bootstraps (Le and Gascuel, 2008; Nguyen *et al.*, 2015; Hoang *et al.*, 2018). Phylogenetic trees were edited in iTOL 4.4 by collapsing branches supported by <500 bootstraps (Letunic and Bork, 2019).

### Synteny network extraction and syntelog clustering

Synteny networks were obtained by extracting the identified HMMERlogs and their syntenic connections from the proteome-wide angiosperm synteny network presented in Zhao and Schranz (2019). Residual non-HMMERlog genes were discarded. Syntenic HMMERlog genes (syntelogs) were clustered with the Infomap algorithm in R (Rosvall and Bergstrom, 2008). Redundant connections were removed, and only syntelogs with a k-core >3 were kept. Synteny networks were visualised and coloured with Cytoscape 3.7.1 (Shannon *et al.*, 2003). Phylogenomic profiles were created by counting the number of syntenic HMMERlog genes per syntelog cluster in all 107 species. Subsequent hierarchical clustering was performed using the heatmap.2 package in R. Synteny information was used to define AP2 subclades and to determine syntenic connections between subclades. To eliminate one-to-many syntenic connections between subclades, a threshold was set at a maximum of 10 connections originating from a single syntelog. Lineage-specificity of synteny was determined by counting the number of syntelog clusters per subclade. Syntelog clusters were considered to be lineage-specific when containing at most two species belonging to other taxa. A cluster was not assigned to a subclade when containing less than 10% HMMERlogs belonging to that particular subclade.

### Cluster overlap index analysis

As a measure for the relative degree to which a certain gene family is conserved, we devised the cluster overlap index. This index is calculated by performing pairwise interspecies comparisons of overlapping syntelog clusters. For example, species *A* has a cluster overlap index of 0.8 with species *B*, when eight out of ten clusters contain syntelogs of both *A* and *B*. Distributions of overlap indices are a measure to compare relative conservation of synteny between *AP2* and *EREBP* subfamilies, i.e. as long as the same input species are used. Angiosperm species lacking syntelogs in both families were excluded from this analysis.

## ACKNOWLEDGEMENTS

We thank Tao Zhao and Setareh Mohammadin for technical assistance and providing scripts. Viola Willemsen is acknowledged for constructive suggestions and ideas for improving the manuscript.

## AUTHOR’S CONTRIBUTIONS

MHLK, KB and MES designed the research. MHLK performed experiments. MHLK and KB analyzed data. MHLK, MES and KB wrote the manuscript. All authors read and approved the final manuscript.

## SUPPORTING INFORMATION

**Figure S1.** Maximum-likelihood phylogenetic tree of AP2 HMMERlogs displaying unfiltered syntenic relationships within and between subclades.

**Figure S2.** Lineage-specific synteny of a *LEC1/L1L* cluster in Brassicaceae.

**Figure S3.** Maximum-likelihood phylogenetic tree of EREBP HMMERlogs displaying syntenic relationships within and between subclades.

**Figure S4.** Copy number variation in *EREBP* subclades in angiosperms.

**Figure S5.** Phylogenomic profile of *EREBP* syntelogs in 107 angiosperm species.

**Figure S6.** Bar plot displaying the number of syntelog clusters and total number of HMMERlogs per *EREBP* subclade.

---

**Figure S1.** Maximum-likelihood phylogenetic tree of AP2 HMMERlogs displaying unfiltered syntenic relationships within and between subclades.

**Figure S2.** Lineage-specific synteny of a *LEC1/L1L* cluster in Brassicaceae. (**A**) Phylogenomic profile of LAFL transcription factor genes across angiosperm genomes. Presence of lineage-specific synteny in Brassicaceae is indicated by a green box. (**B**) Interaction network of LAFL transcription factors showing syntelog clusters per subclade. The Brassicaceae-specific *LEC1*/*L1L* cluster is indicated by a green box.

**Figure S3.** Maximum-likelihood phylogenetic tree of EREBP HMMERlogs displaying syntenic relationships within and between subclades. Syntenic relationships between HMMERlog genes are indicated by grey lines. Subclades are named following the *Arabidopsis* gene nomenclature. Subclades lacking *Arabidopsis* orthologs are annotated with the suffix -L (-like). The *AP2*^*AP2*^ subclade consists of HMMERlogs with only a single AP2 domain, but have higher homology with AP2 HMMERlogs.

**Figure S4.** Copy number variation in *EREBP* subclades in angiosperms. (**A**) Phylogenomic profile of *EREBP* syntelogs across angiosperm genomes. (**B**) Box plot showing the distribution of EREBP HMMERlogs per subclade across angiosperm lineages.

**Figure S5.** Phylogenomic profile of *EREBP* syntelogs in 107 angiosperm species.

**Figure S6.** Bar plot displaying the number of syntelog clusters and total number of HMMERlogs per *EREBP* subclade.

