## Supplementary figures and images for "Phylogenomic analysis of the APETALA2 transcription factor subfamily across angiosperms reveals both deep conservation and lineage-specific patterns"

### Figure S1

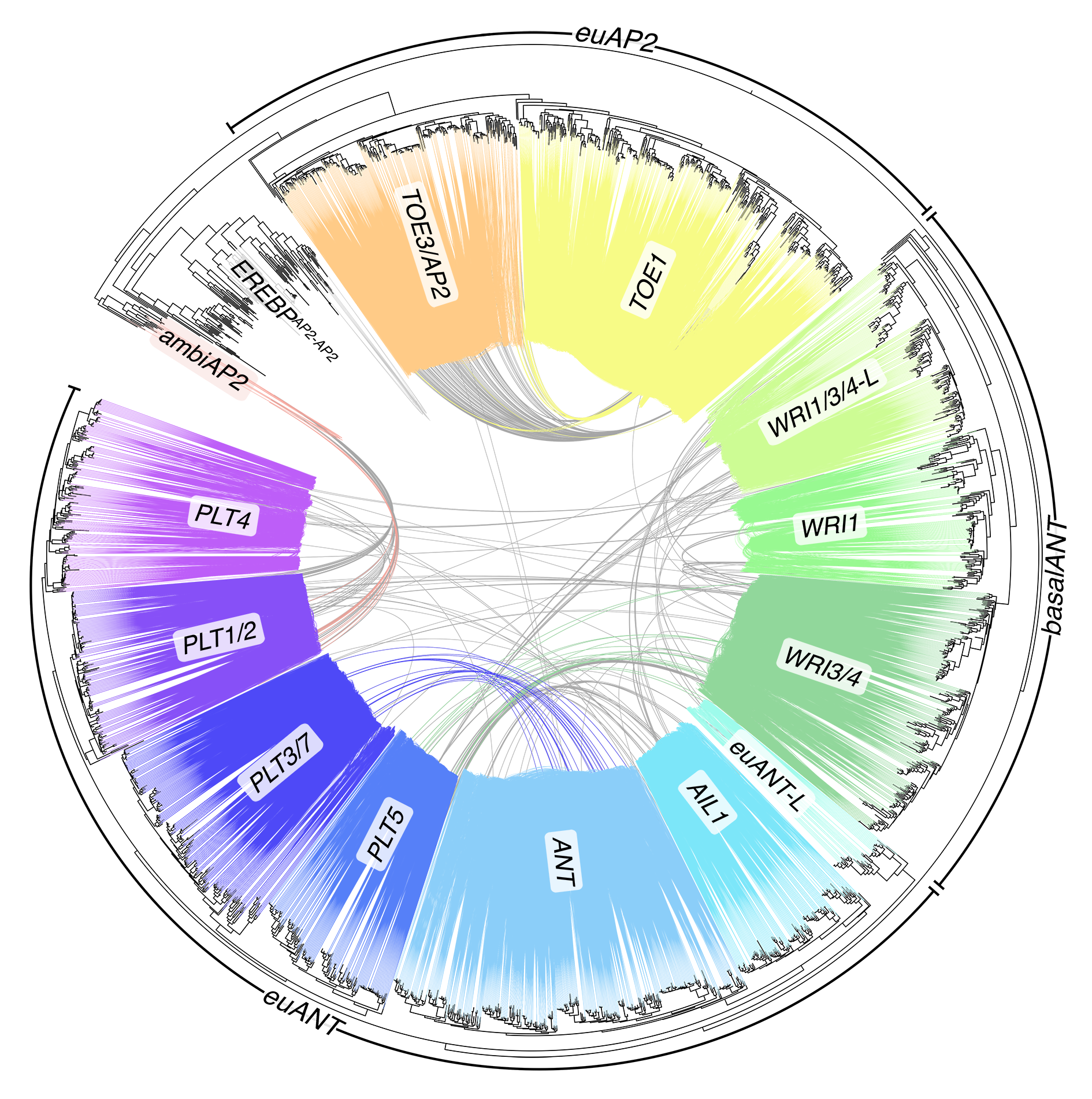

### Figure S2

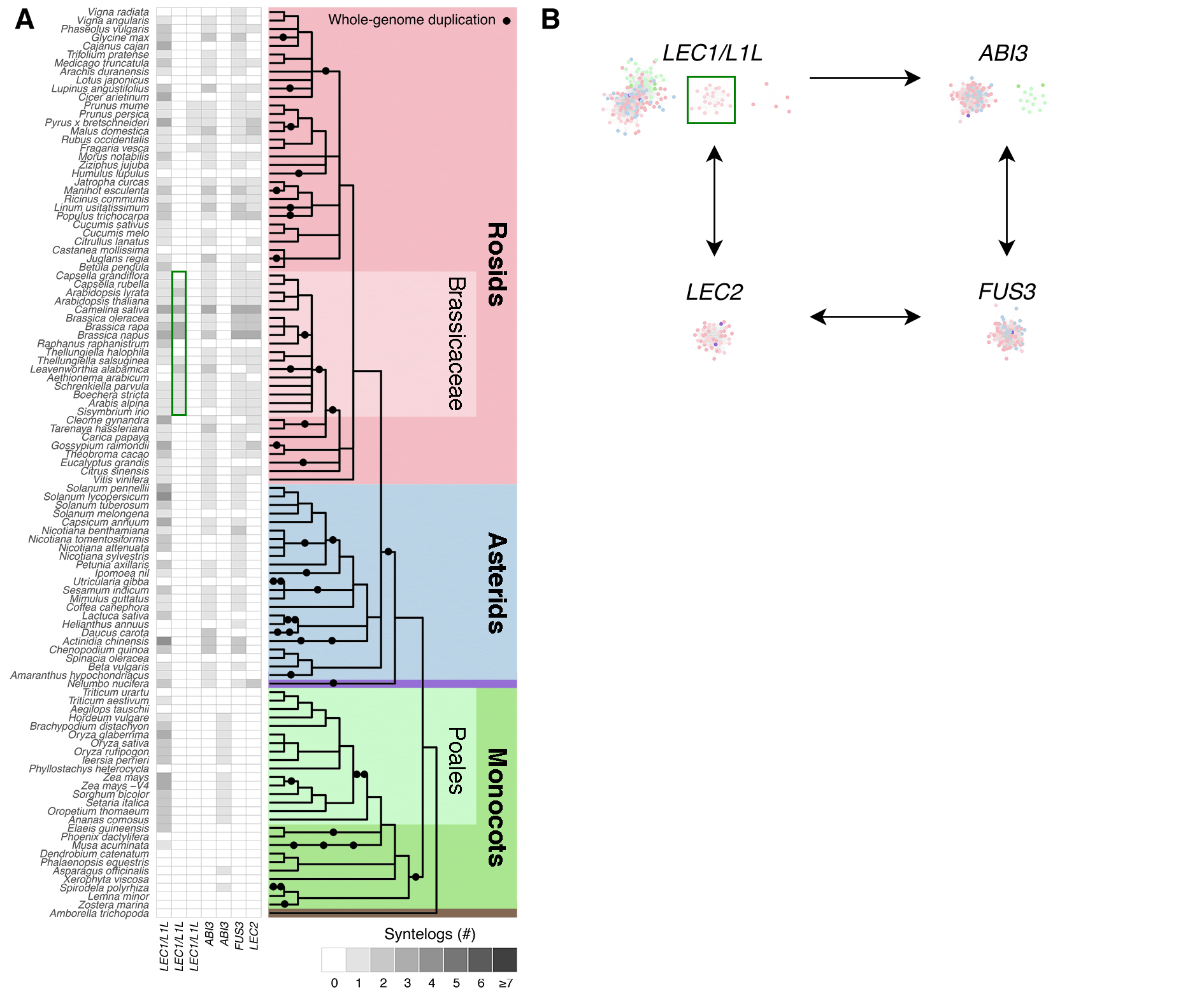

### Figure S3

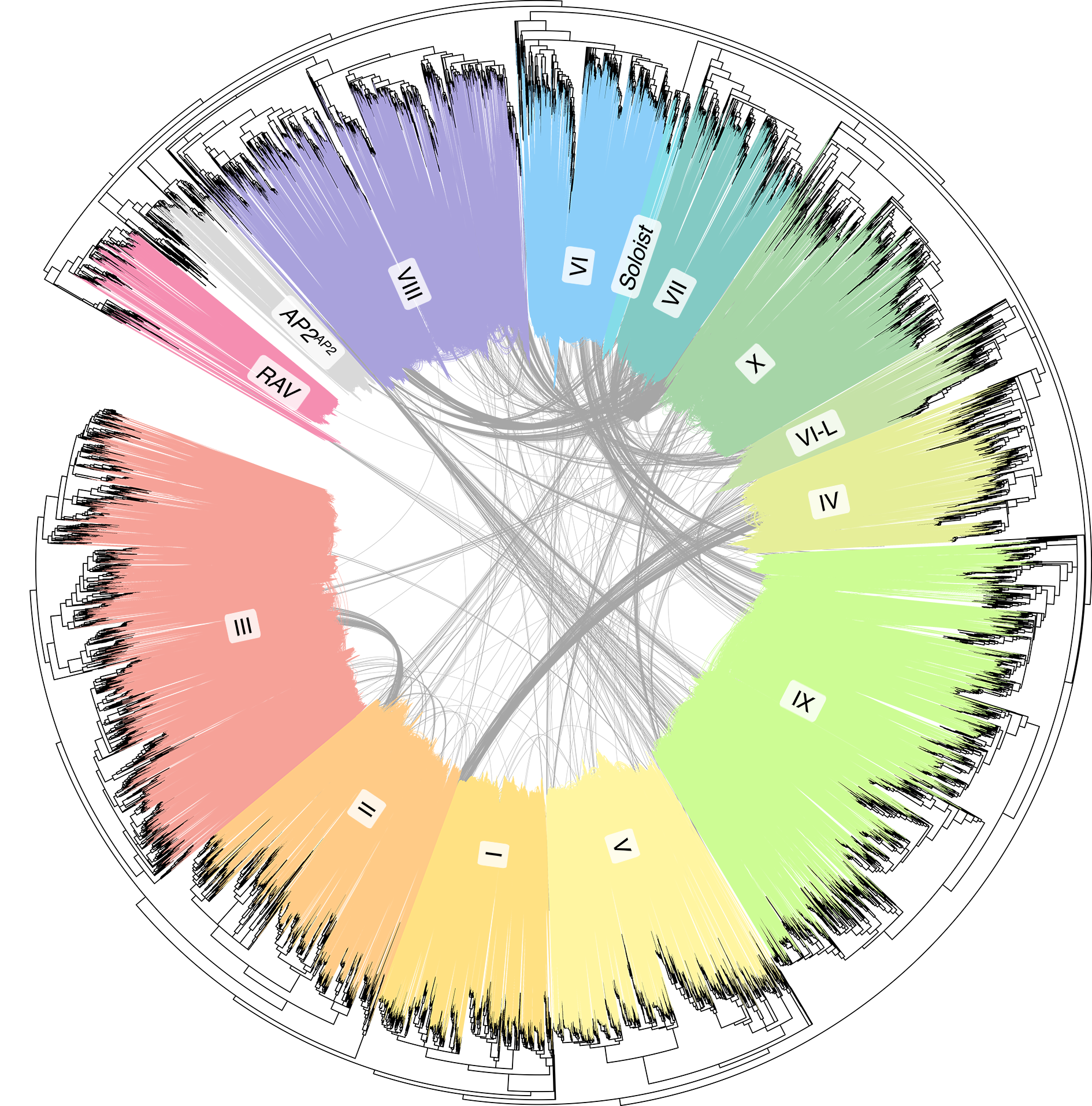

### Figure S4

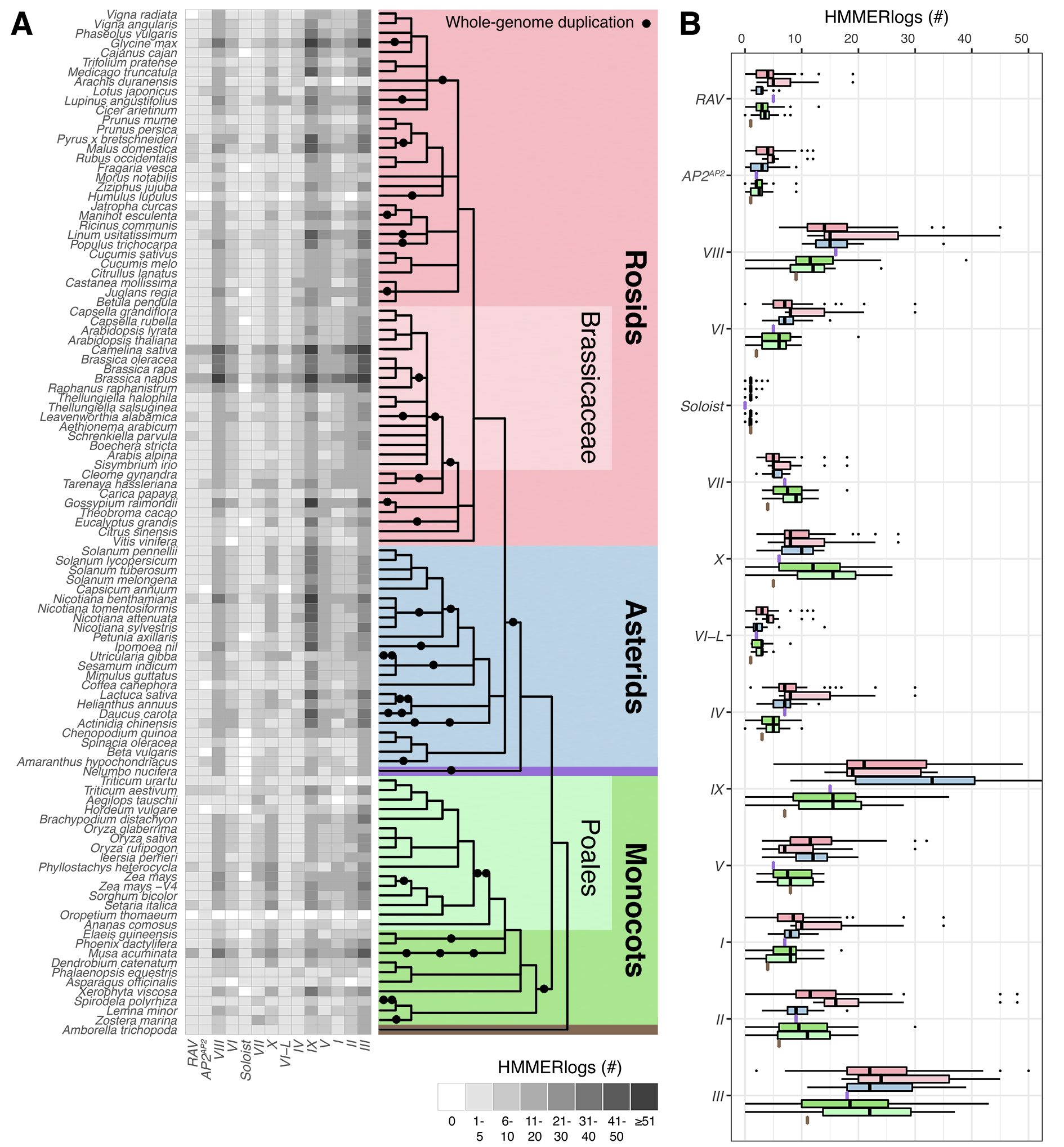

### Figure S5

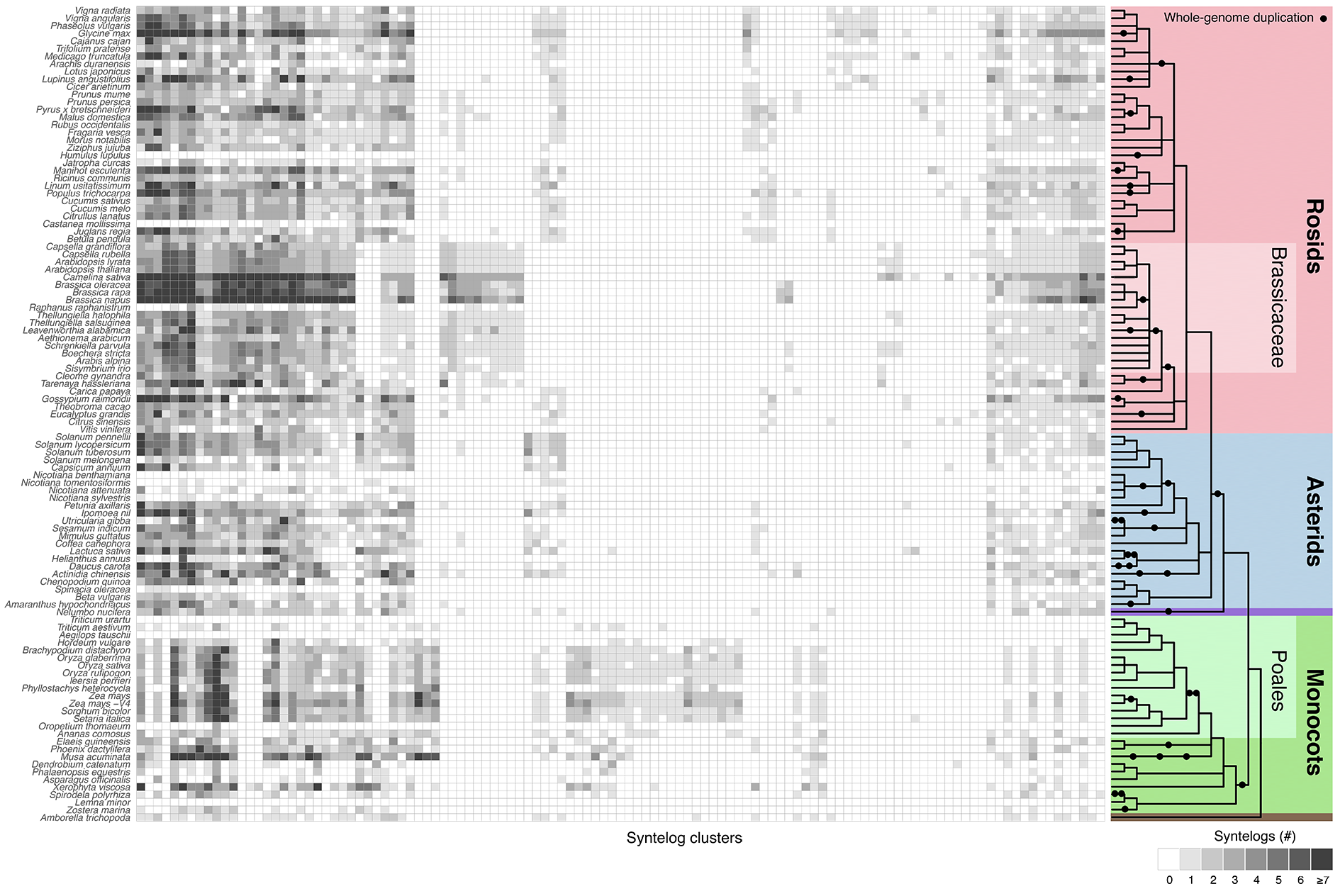

### Figure S6

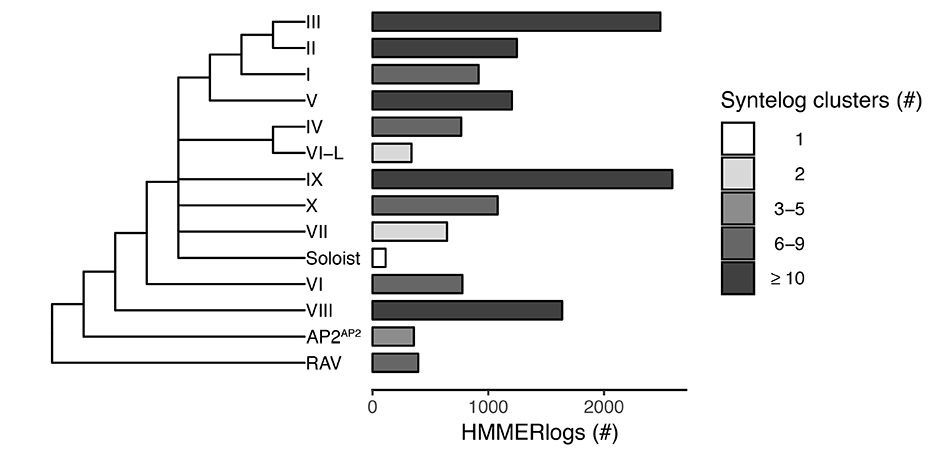
